# *In silico* genomic surveillance by CoVerage predicts and characterizes SARS-CoV-2 Variants of Interest

**DOI:** 10.1101/2024.03.07.583829

**Authors:** K. Norwood, Z.-L. Deng, S. Reimering, G. Robertson, M.-H. Foroughmand-Araabi, S. Goliaei, M. Hölzer, A. C. McHardy

## Abstract

Rapidly evolving viral pathogens such as SARS-CoV-2 continuously accumulate amino acid changes, some of which affect transmissibility, virulence or improve the virus’ ability to escape host immunity. Since the beginning of the pandemic and establishment of SARS-CoV-2 as a human pathogen, multiple lineages with concerning phenotypic alterations, so called Variants of Concern (VOCs), have emerged and risen to predominance. To optimize public health management and to ensure the continued efficacy of vaccines, the early detection of such variants of interest is essential. Therefore, large-scale viral genomic surveillance programs have been initiated worldwide, with data being deposited in public repositories in a timely manner. However, technologies for their continuous interpretation are currently lacking. Here, we describe the CoVerage system (www.sarscoverage.org) for viral genomic surveillance, which continuously predicts and characterizes novel and emerging potential Variants of Interest (pVOIs) from country-wise lineage frequency dynamics together with their antigenic and evolutionary alterations utilizing the GISAID viral genome resource. In a comprehensive assessment of VOIs, VOCs and VUMs identified, we demonstrate how CoVerage can be used to swiftly identify and characterize such variants, with a lead time of almost three months relative to them reaching their maximal abundances. CoVerage can facilitate the timely identification and assessment of future SARS-CoV-2 variants relevant for public health.

## Introduction

In early 2020, infections with a previously unknown coronavirus of probable zoonotic origin in Wuhan, China were first reported^1^. The virus, named SARS-CoV-2, rapidly spread across the globe, causing over 675 million infections and 6.8 million deaths as of March 2023^2^. SARS-CoV-2 has a single-stranded, positive-sense RNA genome with substantial capacity to mutate, reflected in its strain-level diversity of circulating viral lineages and rapid evolution^3,4^. This led to the emergence of several Variants of Concern (VOCs), as designated by the World Health Organization (WHO), with altered phenotypes in transmissibility, virulence, or antigenicity, causing large waves of new infections or reinfections^5–8^.

Generally, viral pathogens such as human influenza and severe acute respiratory syndrome coronavirus 2 (SARS-CoV-2) viruses evolve rapidly, adapting to the human host for efficient replication and spread. Continuous changes on the surface antigens of these viruses allow them to evade host immunity developed through either prior infection from previous strains or from vaccination. This capacity of a virus, known as immune escape, allows the virus to reinfect individuals and, consequently, vaccines protecting against such viruses need to be frequently updated to maintain their effectiveness against currently circulating variants^9^. SARS-CoV-2 in particular demonstrates an increased capacity for immune escape with more recent circulating variants evading vaccine-derived antibodies and convalescent sera^10^. The Omicron variant designated a Variant of Concern (VOC) by the World Health Organization (WHO) in November 2021, and its sublineages BA.2, BA.4 and BA.5 have reduced susceptibility to monoclonal antibodies (mAbs) in clinical use^11,12^. The latest Omicron subvariant, JN.1, which has rapidly risen to predominance in January 2024 with a global prevalence of 72.89%, has over 30 amino acid mutations occurring on the spike protein with an additional mutation L455S and significantly enhanced immune escape compared to its parent lineage BA.2.86^13,14^. This increasing capacity for immune evasion is driven by key mutations throughout the spike protein where they reduce neutralization by antibodies or T-cell based responses^10^.

To monitor SARS-CoV-2 evolution and adaptation as well as enable the timely identification of new VOCs, many countries implemented large-scale viral genomic surveillance programs, leading to the generation of unprecedented amounts of sequence data. As of March 2024, more than 16.5 million sequences were available in the GISAID database^15^. Though web-based platforms offer various analyses based on publicly available sequencing data that support scientists, public health officials, and the general public in making sense of these highly complex data, there remains a need for methods that identify antigenically altered lineages among the numerous circulating lineages, particularly among lineages rapidly rising in frequency, to support public health related decision making in a timely manner. The early identification of such variants is particularly relevant for vaccine updates to ensure continued vaccine efficacy, such as in the case of Omicron, XBB.1.5 and JN.1^16^.

Here, we describe CoVerage (https://sarscoverage.org), an analytics platform that monitors the genetic and antigenic evolution of human SARS-CoV-2 viruses. The platform implements methods that continuously search for pVOIs from global, country-wise viral variant frequency dynamics and predicts relevant evolutionary changes and their antigenic alterations relative to the original Wuhan strain, using publicly available viral surveillance and genome data in a fully automated, timely fashion. The term pVOI, adapted from the WHO defined Variant of Interest (VOI), defines potential Variants of Interest (pVOI) as predicted by CoVerage, which identifies pVOIs as those that increase significantly in frequency over time and rise above a predominance threshold for the first time in a fully automated fashion^17^. We demonstrate the application of the framework for the early detection of circulating Variants of Interest (VOIs), Variants Under Monitoring (VUMs) and VOCs.

## Results

### CoVerage implementation

The CoVerage analytical workflow comprises several stages: (1) input data acquisition and filtration, (2) computational sequence data analyses, (3) creation of template pages based on a bootstrap framework, and (4) visualization in the browser using GitHub Pages (Fig. 1). First, both genomic sequences and the corresponding metadata file are used as inputs for the CoVerage computational analytics pipeline. For this purpose, genomic sequences and the sequence metadata file are downloaded with daily updates from GISAID^18^. Optionally, German sequence data is downloaded from Zenodo, where it is made available by the Robert Koch Institute (RKI) (https://zenodo.org/record/8334829), prior to its submission to GISAID. Case numbers are obtained from the WHO Coronavirus (COVID-19) data repository^19^. The computational workflow performs the analysis of SARS-CoV-2 lineage dynamics by country to identify emerging lineages that may possess a selective advantage, analysis of allele dynamics of the SARS-CoV-2 major surface protein to identify alleles with amino acid changes that may provide a selective advantage, and the *in silico* assessment of antigenic lineage alterations. Antigenic alteration and the lineage dynamics analysis require the GISAID metadata as input, while the spike protein allele dynamics requires both the genomic sequences and associated metadata to infer the coding sequences. Each of these analyses and data downloads are run independently once per week, and the results are updated on the web-server accordingly.

**Fig. 1:**
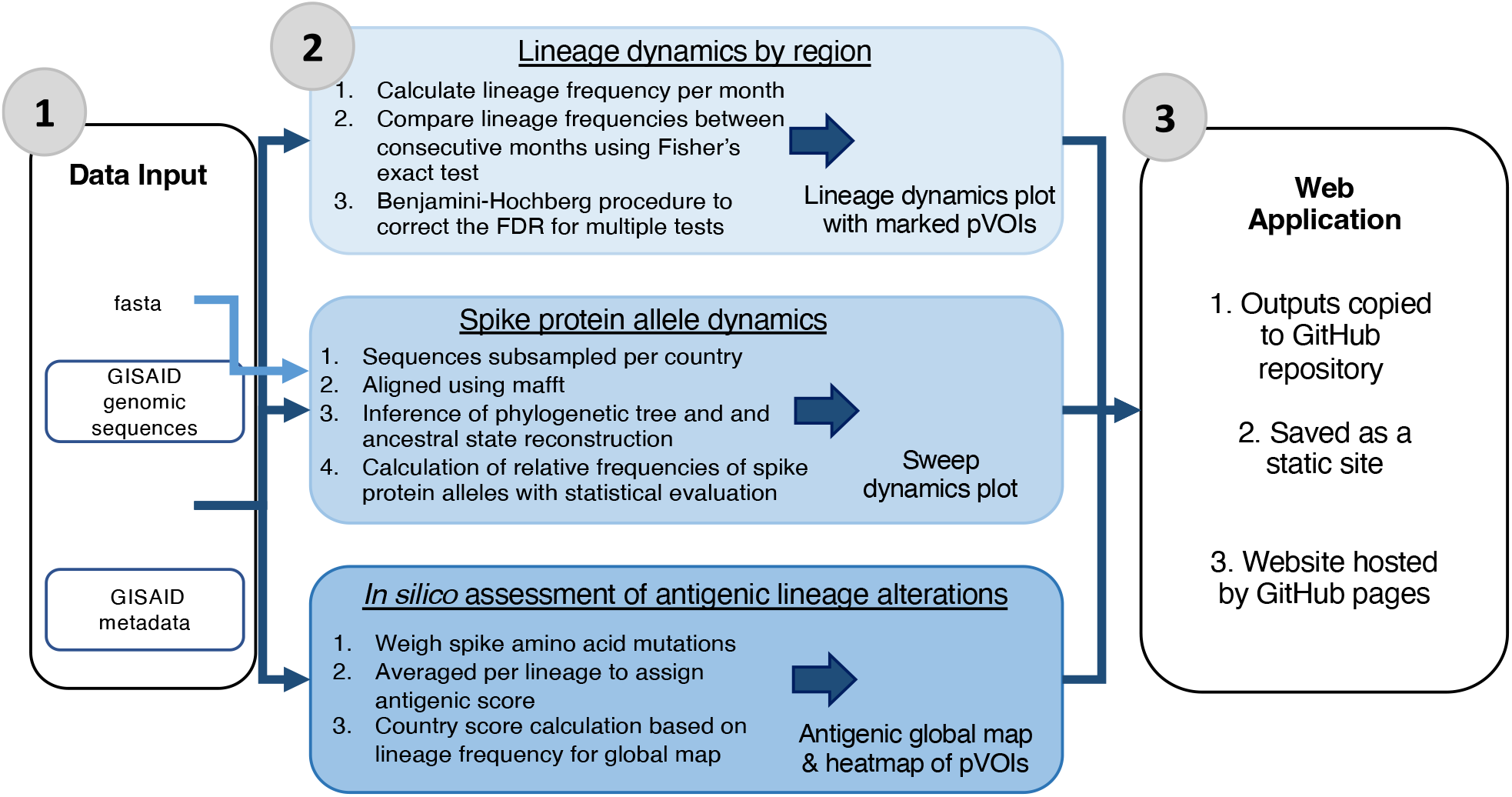
Continuous viral genome analytics provided by the CoVerage platform (sarscoverage.org). Data input includes GISAID genomic sequences and metadata for SARS-CoV-2 isolates from GISAID. These inputs are then fed through three computational workflows, whose outputs are saved to individual GitHub repositories for each analysis. The results are then copied to the main CoVerage GitHub repository, where the individual pages are saved as static sites, which are hosted by GitHub pages. Workflows are run once a week and result pages are updated accordingly.

### Lineage and spike protein dynamics analyses provided by CoVerage

CoVerage provides lineage and spike protein dynamics analyses updated once a week for all countries with sufficient data available (set to more than 2000 sequences) to which one can navigate to via the interactive world map (**Supplementary Fig. 1**). The lineage dynamics analysis suggests potential Variants of Interest for individual countries (**Fig. 1**), as lineages rising in prevalence more rapidly than expected by chance (Methods), which may be due to a selective advantage and increased fitness to spread relative to other circulating viral lineages. Furthermore, CoVerage provides an analysis of spike protein allele dynamics, which identifies “lineage alleles”, corresponding to branches in a viral spike protein genealogy and associated amino acid changes increasing significantly in frequency over time, which can indicate that these confer a selective advantage to the respective lineages^20^ (Methods). For the analysis of lineage dynamics and to identify pVOIs, i.e. lineages that may have a selective advantage to spread in the host population, as well as for identifying sets of amino acid changes on the spike protein rapidly increasing in frequency, we adapted a technique that we developed for recommending vaccine strain updates for the seasonal influenza vaccine^20^ (https://github.com/hzi-bifo/SDplots). There are three methodologies for identifying pVOI lineages: the ‘standard’ method identifies lineages increasing significantly in frequency over time with monthly intervals using Fisher’s exact test. The sub-lineage corrected method for pVOI identification includes genome sequence assignments belonging to sublineages in the pVOI assessment. Finally, the sliding window method uses a windowed time period for the analysis to achieve a more fine-grained analysis of shorter time intervals (Methods).

For the lineage dynamics analyses, Pango lineages were used as nomenclature, which define epidemiologically relevant phylogenetic clusters, in which new lineages are only designated if the lineage has high coverage and contains a sufficient number of sequences^21^. All methods utilize Fisher’s exact test to identify lineages that are significantly increasing in frequency over two consecutive time intervals and correct for multiple testing using the false discovery rate (FDR, α = 0.05)^22^. This identifies pVOIs as lineages that are both significantly on the rise and increase above a predominance threshold of 0.1 within the same time frame. Notably, lineages may also be falsely identified as pVOIs due to unrepresentative sampling, e.g., data biases towards certain areas, large clonal outbreaks, or population bottlenecks. Therefore, identified pVOIs should be evaluated carefully, e.g. in combination with epidemiological data and experimental evidence showing that amino acid changes and altered positions observed in a pVOI lineage are likely to confer a selective advantage.

### *In silico* assessment of antigenic lineage alterations

CoVerage identifies antigenically altered lineages using a novel method which scores evolutionary changes based on an antigenic alteration matrix (Methods). This matrix is based on genotype-to-antigenic phenotype associations observed in the long-term evolution of seasonal influenza A (H3N2) viruses (**Fig. 1**). Amino acid changes in the spike protein are weighed as the spike protein plays a critical role in SARS-CoV-2’s binding to host receptor cells and is a key target for vaccines and antibody treatments^23^. These can be visualized in a monthly map of the global depicting antigenically altered circulating lineages or as a heatmap with antigenically altered lineages and their corresponding frequency. Significantly altered lineages are denoted with an asterisk, otherwise altered lineages are ranked by frequency. To identify significantly antigenically altered lineages, a z-score standardization is applied to the circulating lineages of the month, where lineages with a z-score greater than one, or greater than one standard deviation from the mean, are denoted as altered compared to the other circulating lineages for that month. These lineages are then visualized in the antigenic alteration heatmap on the CoVerage platform.

### Widely spreading antigenically altered SARS-CoV-2 lineages can be detected early

For validation of the pVOI identification and antigenically altered scoring methodologies created by CoVerage, we determined, for pVOIs and antigenically altered lineages predicted until the end of June 2024, the time it took for these lineages to reach their peak frequencies (**Fig. 2**). The frequency for each lineage in each country was calculated using a one week sliding window with a step size of two days. From here, the peak sequence count, the corresponding frequency, and the date of the peak were identified. To ensure robustness and mitigate bias from rare lineages or inadequate sequence sampling, we included only those records where the peak sequence count was at least 5 within the respective window. In our comprehensive evaluation, a total of 404 detected pVOIs from 91 countries were analyzed, corresponding to 1360 pVOI predictions for individual countries. On average, CoVerage identified the pVOI using data from 49 days earlier than the date when it reached its peak frequency. For the countries with more data available (peak sequence count >=100 for the respective lineage), this lead time extended to 78 days (**Fig. 2**). The antigenically altered pVOIs have a day difference of 72 compared to 24 for non-altered pVOIs and reached a much higher peak frequency than the non-altered ones (**Fig. 2a**), indicating the relevance of antigenic alterations for providing a selective advantage to variant lineages.

**Fig. 2:**
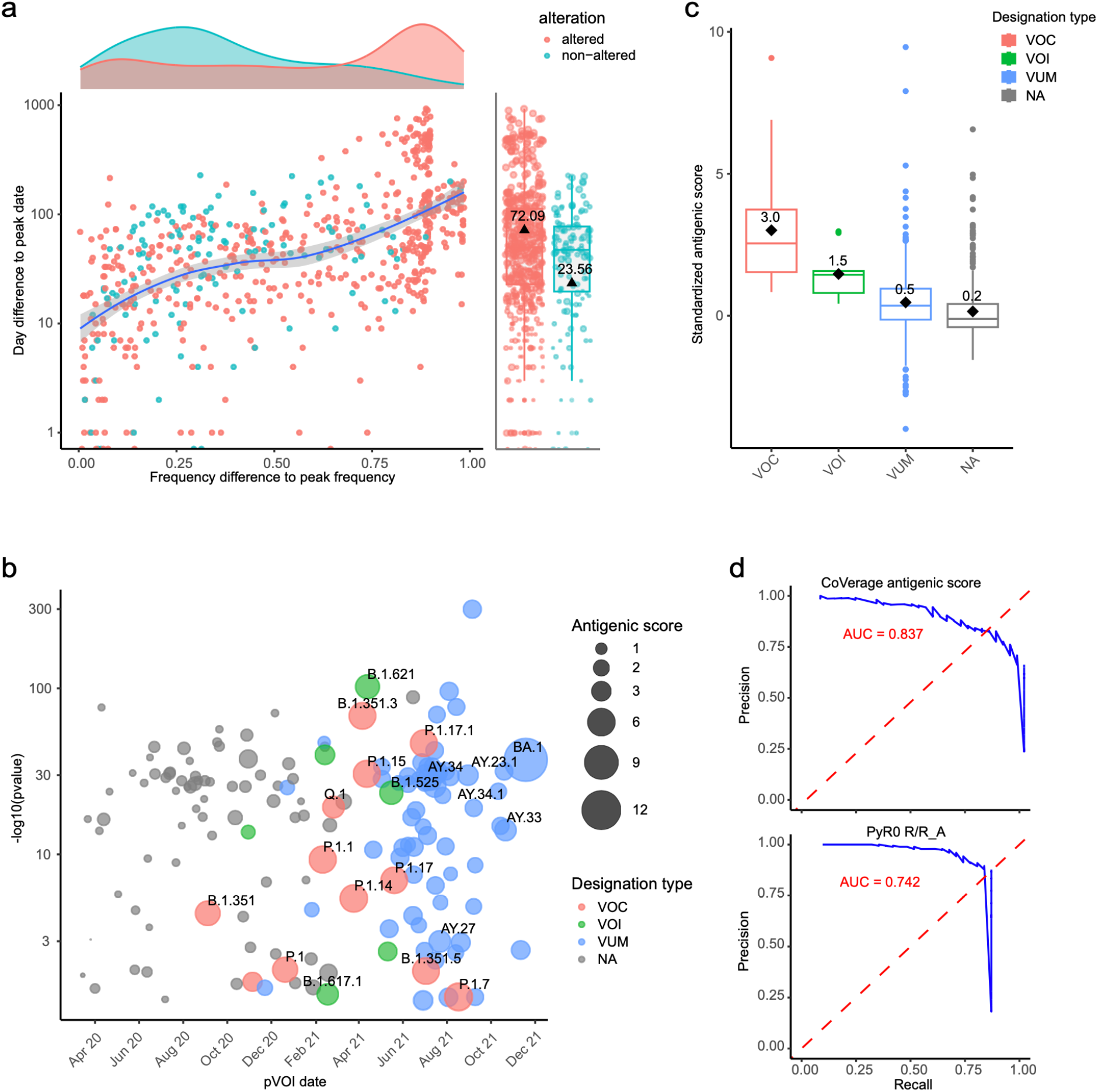
Performance evaluation of CoVerage. **a** The difference in days and frequency of pVOI identification of lineages relative to their peak frequencies. The dots in red indicate predicted antigenically altered pVOIs, while the turquoise dots represent predicted non-altered pVOIs. **b** Antigenic scores of pVOIs identified using data before December 2021. Colors represent different WHO designations. The x-axis shows the month of identification, with “Apr. 20” indicating April 2020. The y-axis displays the -log10 transformed p-value of pVOI detection. The size of the points represents the antigenic score of the pVOI. **c** Standardized (z-score) antigenic scores for variants and their sublineages for different WHO variant categories designations until June 2024. Here the horizontal bar within the box indicates the median, while the black diamond and the number above it represents the mean. “NA” indicates pVOIs not designated by WHO. **d** PRAUC of CoVerage antigenic score and PyR0 R/R_A_ in classifying/predicting lineages as WHO-designated or non-WHO- designated VUMs, VOIs and VOCs. Data for both methods were collected before December 2021, with the ground truth including WHO-designated lineages up to May 2022 to demonstrate early detection capabilities. In **b** and **c**, “NA” indicates pVOIs not designated by WHO.

Overall, 320 pVOI lineages identified by CoVerage belong to the 45 officially designated VOCs, VOIs, and VUMs, covering 33 of these designated lineages, resulting in a precision of 79% (320/404) and a recall of 73% (33/45). Among the 12 lineages not captured by pVOIs, 10 are VUMs, thus the lowest category of relevance from public health monitoring. No VOCs were missed. On average, these pVOIs were first identified using data collected 84 days before their official WHO designation.The pVOIs not designated by WHO were primarily identified in 2020, during the early stages of the pandemic, when available sequence data was oftentimes limited and less from systematic representative surveillance efforts, as in the following years (**Fig. 2b**). Notably, the antigenic scores and standardized scores of pVOIs that were also WHO designated lineages were much higher than those of the non-designated pVOIs, and increasing in order of VUMs, VOIs and VOCs, in line with their proven relevance for public health decision making (**Fig. 2c**). To evaluate the benefits of our method relative to a sequence analysis/based method, we compared PyR0^24^ relative to the antigenic scoring method (**Fig. 2d**), which showed that the antigenic alteration score of variants of CoVerage predicted VOCs, VOIs, and VUMs more accurately than the R/R_A_ score used by PyR0, with an Area Under the Precision-Recall Curve (PRAUC) of 0.84 vs 0.74, respectively. Overall, these findings underscore our method’s capability to effectively predict the ascent of lineages relevant for public health decision making with a growth advantage well ahead of their peak frequency, providing critical lead time for public health interventions.

### Case study 1a: CoVerage lineage dynamics allow for timely identification of Omicron as pVOI

We retrospectively analyzed data from the time of emergence of the Omicron lineage or Pango lineage BA.1 as the Omicron lineage, to assess the detection of pVOIs, prediction of antigenic alterations (Case study 1b), and of relevant amino acid changes (Case study 1c). This was after finalizing all methodological settings of pVOI detection in first year of the pandemic, and utilizing an antigenic alteration scoring method, which scores all amino acid changes in the spike protein relative to the original Wuhan strain, thus not including knowledge about relevant sites for Omicron or other variants. Using viral genome information provided until November 23rd of 2021, the day when the first Omicron sequences from South Africa identified under viral genomic surveillance^25^ were submitted, the CoVerage lineage dynamic analysis suggested BA.1 as a pVOI, which then occurred with relative frequency of 0.27 among South African isolates (**Fig. 3a**). The detection of BA.1 as a pVOI was only three days prior to the swift designation of Omicron as a VOC by the WHO^26^, owing to the highly efficient work and data release of South-African scientists. The isolates used in the analysis were sampled between November 14th and November 16th, demonstrating how the methodology can rapidly identify novel lineages of concern when data is submitted to the GISAID database in a timely manner (**Fig. 3**). The global heatmap visualizing the relative frequencies of all pVOIs detected by CoVerage lineage dynamics analysis in countries worldwide, Omicron (BA.1) was identified in November 2021 in South Africa (**Fig. 3c**), and then rose rapidly in frequency together with multiple sublineages such as BA.1.1, BA.1.15 and BA.1.17 to the most prevalent lineage worldwide by December 2021 (**Fig. 3d**).

**Fig. 3:**
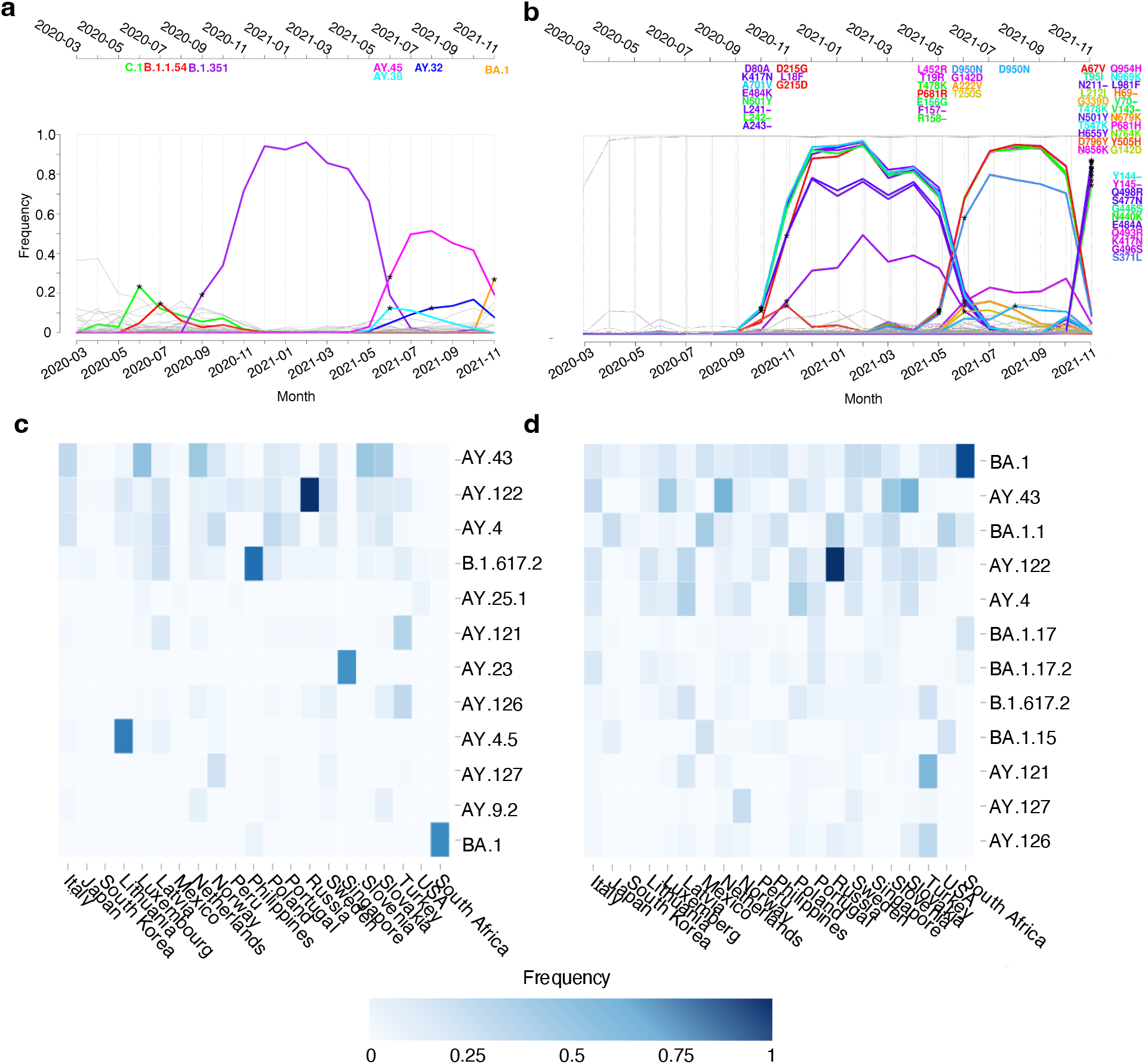
a Lineage dynamics plot for Pango lineages of SARS-CoV-2 genomes from South Africa submitted to GISAID by November 23rd, 2021. Colored lines represent identified pVOIs with their Pango lineage names given above. Asterisks indicate the month in which they were identified as being significantly on the rise and increasing in predominance above a predefined threshold of 0.1. All other lineages are shown in gray. **b** SD plot for South Africa on spike protein sequences available until the end of November 2021. Similar to the lineage dynamics plot, the asterisks represent when a spike protein allele with certain amino acid changes significantly rises in frequency and the color of the curve corresponds to the associated amino acid change. Amino acid changes are specified at the top of the plot, based on the time point when they were identified as significant. The asterisks in November 2021 indicate the detection of the emerging Omicron lineage (officially announced as a VOC on November 26th, 2021)^28^, in May 2021 of Delta (Pango lineage B.1.617.2, detected with CoVerage first as a pVOI for India in March of 2021 and announced as a VOC on May 11th, 2021)^9^, and Beta (B.1.351, announced as a VOC on November 29th in 2020) in September of 2020^29^. Asterisks in November 2020 indicate within lineage variation of Beta, in June and August of 2021 of sublineages AY.45, AY.38, and AY.32 of the Delta variant, respectively. Relative frequencies for SARS-CoV-2 pVOIs identified in individual countries and most abundant worldwide. **c** November 2021 and **d** December 2021. The color scale ranges from dark blue, which indicates a frequency of one, to white, indicating a frequency of zero. The top 50 countries with the highest lineage frequencies are shown.

Earlier, CoVerage identified Beta (B.1.351) in September of 2020 and several Delta sublineages (AY.45, AY.38 and AY.32) throughout June and August of 2021 in South Africa as pVOIs (**Fig. 3a**), which were later designated VOCs by the WHO^27^. Furthermore, lineages C.1 and B.1.1.54 were identified in June and July of 2020 in the standard and sublineage-corrected analyses, which rapidly increased in frequency among sequenced isolates from South Africa in the month of their designation and by August 2020, the C.1 lineage was the most geographically widespread lineage in South Africa^25^.

### Case study 1b: Spike protein allele dynamics suggest emerging lineages and their associated amino acid changes in South Africa

In the spike protein allele dynamics, for South Africa, CoVerage detected the Beta lineage via a set of associated changes in the spike protein in October 2020, as well as Delta with its amino acid changes in May 2021 (**Fig. 3b**). Key identified substitutions of the Beta lineage include the amino acid deletion at position 241-243, which is associated with reduced binding of certain nAbs, and K417N, that was also present in Alpha^30^. K417N, in combination with the E484K change in Beta, has demonstrated a reduced mAb neutralization, but does not display this capacity alone^11^. Key amino acid changes known to affect the phenotype of the Delta lineage identified in allele dynamics include L452R, which has previously been shown to be an important driver of adaptive evolution; T478K, which facilitates improved immune escape; and P681R, which enhances furin cleavage^30,31^. At the time of the emergence of BA.1, a lineage allele including all 35 amino acid changes representative of BA.1 in November 2021 (**Fig. 3b**). Among the changes known to alter the phenotype and provide a selective advantage are N501Y, which increases the binding affinity to the host angiotensin converting enzyme 2 (ACE2) receptor cells, E484A, which is a site relevant for immune escape, and P681H, which enhances spike cleavage^30,31^. Due to the extensive divergence of Omicron from other circulating variants, further sampling of more related lineages would have been needed to resolve changes to the ones most likely to providing selective advantages only^32^. Taken together, the results demonstrate how the CoVerage allele dynamics analysis can be used to identify emerging VOCs and their distinctive amino acid alterations based on the ecological dynamics of the spike protein allele in the viral epidemic, without their formal classification as novel lineages.

### Case study 1c: CoVerage predicts antigenic alterations for emerging Omicron variant

CoVerage provides an antigenic scoring analysis that identifies antigenically altered variants based on changes in the SARS-CoV-2 spike protein. Lineages above the selected standardized score threshold and their relative frequencies are shown as a monthly updated heatmap online (**Fig. 4**). In November 2021, Omicron (BA.1) was assigned an antigenic alteration score of 14.47 (z-score of 7.91), more than three times that of Delta (B.1.617.2; 2.11 with a z-score of -0.28) and one and a half times the antigenic score of Gamma (P.1; 5.22 for November 2021, no z-score was assigned as it did not meet the frequency threshold that month), in line with its more pronounced antigenic change^33^. Since December 2021 and continuing into January 2022, Omicron sublineages circulated globally at higher frequencies (**Fig. 3**). Some of these lineages, BA.1.2 and BA.1.21 had even higher antigenic scores than BA.1 in December (15.05 with a z-score of 1.68 and 15.93 with a z-score of 1.84, respectively). The rapid spread of Omicron and its sublineages can also be observed from the global antigenic change maps for October, November, and December 2021 (**Fig. 4**, Methods), which reflect the antigenic alteration scores weighted by their corresponding frequencies per country. For South Africa, in November 2021 the country’s antigenic score rose to 12.86 from 2.40 in the previous month (**Fig. 4b**). Subsequently in December, country antigenic scores increased globally as Omicron and its sublineages rose in frequency, replacing previously circulating lineages (**Fig. 4c**).

**Fig. 4:**
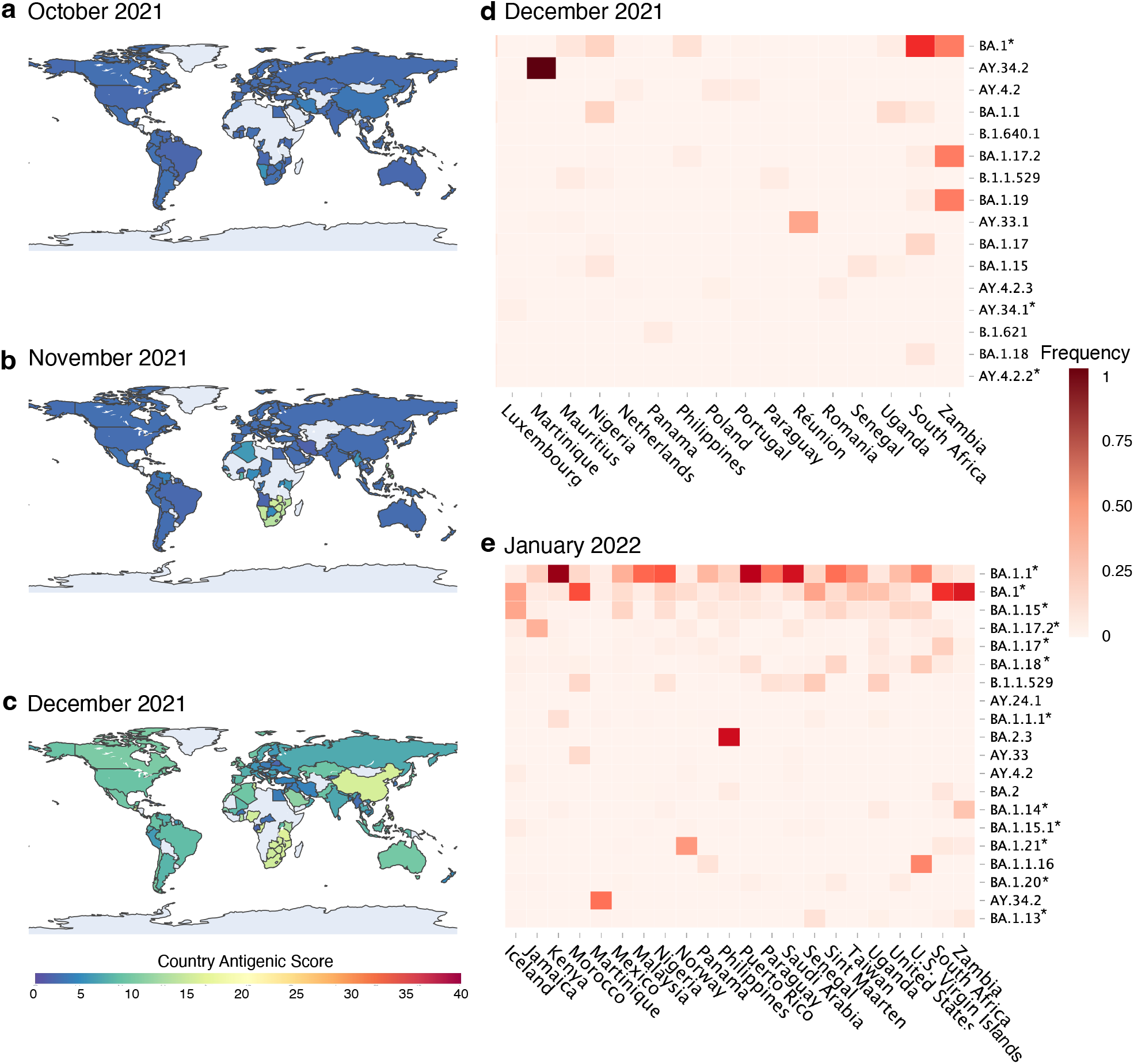
Global map depicting the assigned antigenic alteration score per country over the course of three months when Omicron became a predominant circulating VOC: a October 2021, b November 2021, and c December 2021. A higher score (warmer colors) means that antigenically altered lineages circulate with higher frequency in that country. Dark blue represents an antigenic score per country of 0, while red depicts an antigenic country antigenic score of 40. Heatmap with lineages predicted to have substantial antigenic alteration relative to Wuhan-HU-1 and their frequencies per country as of **d** December 2021 and **e** January 2022. The countries with the highest frequencies are shown and the antigenically altered lineages are ordered from highest to lowest frequency. The dark red color indicates a frequency of 1, while the lighter color red represents a lower frequency closer to 0.

### Case study 2: Screening with CoVerage for antigenically altered pVOIs in 2023

When analyzing available data until March 31, 2023 with CoVerage (Supplementary Tables 3-5), 48 lineages were identified as pVOIs for January, 46 lineages in February, and 37 lineages in March. Most of these are sublineages of BQ.1 and XBB, denoted as BQ.1* and XBB* as per WHO specification^27^. Using CoVerage, the BQ.1 lineage, which was first identified in Nigeria^34^, was detected as a pVOI also in Nigeria in August 2022 (**Fig. 5a**) and designated as a Variant of Interest by the WHO on September 21st, 2022^35^. In the final quarter of 2022 and in the beginning of 2023, BQ.1* was one of the major circulating lineages globally, with BQ.1 and the sublineage BQ.1.1 together covering over 50 percent of submitted sequences in January 2023^36^. Of the XBB sublineages, XBB.1.16 was identified by CoVerage as a pVOI in India in February 2023, two months prior to its VOI designation on April 17, 2023 (**Fig. 5b**)^27^. XBB.1.5 was also identified as significantly antigenically altered in November 2022, with an alteration score of 19.34 (z-score of 1.78)^37^, prior to its VOI and pVOI designations in January 2023 (**Fig. 5c**). BQ.1 and sublineages were quickly overtaken by the XBB.1.5 lineage, which in March 2023 represented over 50% of global sequences^38^. In the antigenic alteration maps, the change from BQ.1.1 to XBB.1.5 as one of the major circulating lineages is evident from January, February, and March 2023 (**Fig. 5d,e,f**). Finally, the WHO recommended an update of COVID-19 vaccine formulations to include the XBB.1.5 lineage in May 2023^39^.

**Fig. 5:**
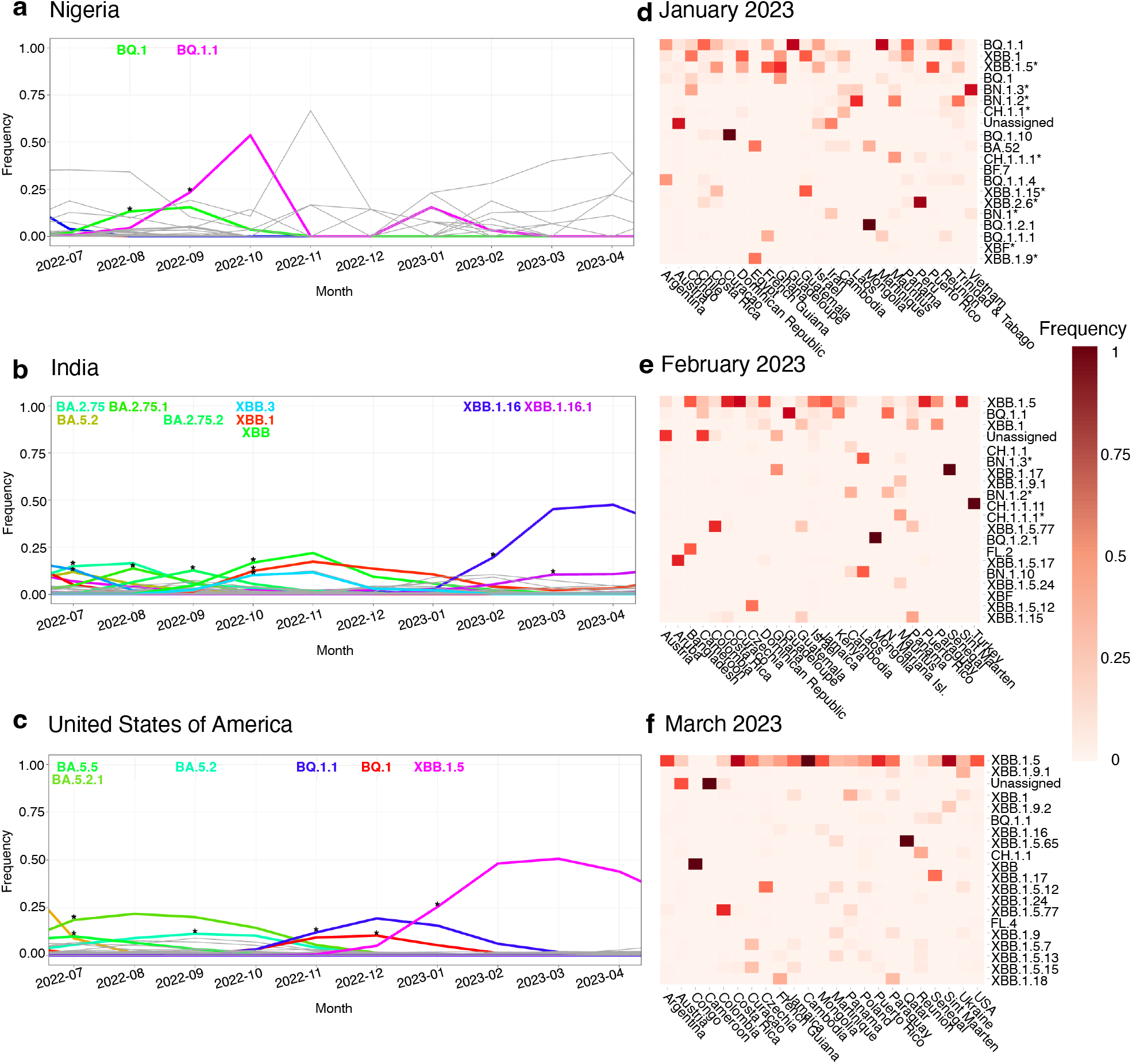
Country-wise lineage dynamics plots for a Nigeria, including results from July 2022 to April 2023, b. India, including results from July 2022 to April 2023, and c the United States, including results from July 2022 to April 2023. **a** The BQ.1 lineage is shown in green and the BQ.1.1 lineage in pink. The asterisks represent a significant rise in frequency, which is seen in August 2022 for BQ.1. **b** The XBB lineage is shown in bright green (October 2022) and the XBB.1.16 lineage is shown in dark blue, XBB.1 in red, XBB.3 in light blue, and BA lineages are shown in shades of green (BA.2.75 and BA.5.2 in July 2022, BA.2.75.1 in August 2022, BA.2.72.3 in September 2022). **c** BA lineages are shown in shades of green and light blue (BA.5.5 and BA.5.2.1 in July 2022, BA.5.2 in September 2022), BQ.1.1 in purple, BQ.1 in red, and XBB.1.5 in pink. The asterisk represents a significant rise in frequency for that lineage. Heatmaps with antigenically altered lineages and their relative frequencies per country in **d** January 2023 **e** February 2023, and **f** March 2023. The lineages are ordered from highest to lowest frequency and the dark red color indicates a frequency of 1, while the lighter color red represents a lower frequency closer to 0.

The increased antigenic alteration scores of the Omicron BQ* and XBB* lineages align with their demonstrated capacity for immune evasion, with their reduced neutralization by select mAbs^36^. Key amino acid changes on the spike protein facilitate immune escape, most notably F486P on the receptor binding domain of XBB.1.5^40^ (**Supplementary Fig. 2**), which also increases infectivity by improving the binding of the spike protein to host ACE2 receptor cells^41^, e.g. in the United States. Furthermore, the N460K mutation of BQ.1.1 has been associated with the loss of neutralizing activity of NTD-SD2 and class I mAbs, while the K444T and R346T changes in BQ.1.1 may also impair the potency of class III mAbs^36^.

Towards the end of 2023, the JN.1 lineage, a BA.2.86 descendant, was designated a Variant of Interest by the WHO on December 18, 2023^42^, two months after its demarcation as a pVOI in Portugal via the lineage dynamics analysis. One of the key amino acid changes is L455S^43^, which was identified e.g. via the allele dynamics analysis in December 2023 in Germany, along with the L157S, N450D, L452W, and N481K changes. Both the N450D and the L452W amino acid changes on the receptor binding domain (RBD) of the spike protein improve viral evasion from multiple mAbs^44^. In experiments, JN.1 showed more extensive resistance to RBD class 1, 2, and 3 antibodies as well as to monovalent XBB.1.5 vaccine sera compared to BA.2.86, which is also reflected in its increased antigenic score for December 2023 of 23.65 (z-score of 0.59) versus 21.33 (z-score of -0.48), respectively^43,4543^. In late April, the WHO advised that future formulations of the COVID-19 vaccines should focus on the JN.1 strain^16^. Subsequently, due to the continued evolution of SARS-CoV-2, the recommendation for the preferred strain was changed to the KP.2 sublineage, if feasible^46^.

## Discussion

Due to the ongoing and rapid genetic and antigenic evolution of circulating SARS-CoV-2 viruses, detecting emerging Variants of Concern that are on the rise to predominance as early as possible is crucial for public health decision making. This included nonpharmaceutical interventions in the early stages of the pandemic and more recently, updating the vaccine composition to ensure continued effectiveness^16,39^. Here, we describe CoVerage, a genomic surveillance platform that continuously monitors globally provided SARS-CoV-2 genomics data, to identify and characterize potential Variants of Interest from the circulating SARS-CoV-2 lineage diversity. CoVerage implements three types of innovative methods for this purpose: (a) a method for the *de novo* detection of potential Variants of Interest that may spread more efficiently than others, (b) a method to identify amino acid changes in the major surface spike protein that may confer a selective advantage, and (c) a method for scoring the degree of antigenic alteration of individual sequences, lineages and the circulating viral diversity per country. To ensure maximal relevance for viral surveillance, CoVerage provides up-to-date predictions of current pVOIs and their antigenic alterations for all countries around the globe with sufficient data available once a week.

Principally, all pVOIs identified in an individual country in one or more countries within the last month are detected and reported by CoVerage in the global pVOI heatmaps. A systematic assessment of these pVOI predictions in combinations with their antigenic alteration predictions showed that this accurately identified 89% of the VOIs and VOCs designated by the WHO since the establishment of SARS-CoV-2 in the human population on average almost three months before their peak abundances were reached. If including also VUMs, a precision of 79% and recall of 73% was thus reached via the pVOI predictions, and CoVerage antigenic alterations scores in ROC analyses also demonstrated high predictive value. The pVOIs and antigenically altered variants identified by CoVerage thus comprise a somewhat larger, but highly informative set of variants with a potential selective advantage for identifying the VOIs, VOCs and VUMs defined by the WHO. The latter are required to show early or confirmed signs of a growth rate advantage, together with further criteria such as the presence of genetic changes that are suspected, predicted or known to affect virus characteristics, along with increasing case numbers or other apparent epidemiological impacts indicating an increasing public health risk, as confirmed by a panel of experts, while the latter contains genetic changes suspected to affect viral characteristics with early signals of growth advantage but either phenotypic or epidemiological impact remains unclear^47^. CoVerage predictions are fully reproducible as they are derived from a defined set of input data, with a fully deterministic, statistical assessment of globally available data, to support and facilitate further expert assessments.

Notably, CoVerage’s detection depends on the extent and quality of ongoing viral genomic surveillance programs for individual countries, as the analysis is done in a country-wise manner and may also be affected by population genetic effects^48^, such as population bottlenecks, when case numbers are low, or travel restrictions between countries are in place, such as in early phases of the pandemic. In terms of data, CoVerage draws on the international GISAID data resource and combines it with other repositories where data is available in advance, such as the genome and metadata published on GitHub/Zenodo by the German Robert Koch Institute, to decrease the time to detecting new, relevant variants^49^. Detection may be affected if genomic surveillance would be decreased further in the future with the virus becoming endemic.

In several case studies, we show how CoVerage detected relevant VOCs as pVOIs along with their antigenic alterations in a timely manner and tracked their ongoing, global spread. In November 2021, Omicron was identified as a pVOI and assigned a substantially increased antigenic score compared to previously circulating lineages such as Delta and Gamma, which was evident also in the country-wide score for South Africa. Further identified VOIs included Omicron’s initial sublineages, BA.1 and BA.2, in December of 2021 and January of 2022, respectively. Consistent with the rapid designation of Omicron as a VOC by the WHO, which in addition to sequencing information also considers epidemiological evidence, CoVerage identified this pVOI from the submitted data just three days before. In early 2023, the majority of pVOIs identified by CoVerage were sublineages of BQ.1 and XBB, both of which have demonstrated improved immune escape due to key mutations throughout the spike protein^36^, many of which could also be found in the spike protein allele dynamics plots of individual countries. By May of 2023 the WHO recommended updating vaccines to match XBB.1.5, which had been identified as a significantly antigenically altered lineage in November 2022 by CoVerage, and subsequently, updated vaccine recommendations to match the circulating JN.1 in April 2024, which had also been identified as an antigenically altered pVOI in October 2023^16,39^. Altogether, this shows the combination of analyses provided by CoVerage facilitates the timely detection and characterization of relevant SARS-CoV-2 lineages for public health concerns.

CoVerage predicts antigenically altered lineages with a novel method that scores evolutionary changes using an antigenic alteration matrix defined from genotype-to-antigenic phenotype association mappings from long-term evolution of seasonal influenza A (H3N2) viruses. The method assesses all changes in the spike protein, which is key immunogen and protein for the binding of SARS-CoV-2 to host receptor cells, and as such is a target for vaccines and antibody therapy^23^. We show that this methods in combination with pVOI predictions allows to confidently identify priority variants and their sublineages over the past couple of years, that were defined by the WHO of public health relevance, such as VOCs, VOIs and VUMs, with on average almost three months prior of them reaching their maximum global frequencies. In evaluating antigenic alteration predictions, we found that predicted antigenic alteration scores are reflective of how well the evolutionary changes affect the neutralization achieved by antibodies. Studying antigenic scores, acquired mutations and their positioning on the 3D structure could thus reveal further insights into the mechanistic basis of varying viral variant neutralization.

Taken together, CoVerage is a unique web-based resource to identify potential Variants of Interest of SARS-CoV-2 in a timely manner, along with suggesting their degree of antigenic alteration, and alleles of the major surface protein with specific amino acid changes that may provide a selective advantage. Notably, there are other web-based resources to track SARS-CoV-2 variants, among other viruses, and their viral fitness and evolution. NextStrain, for instance, not only established a viral lineage nomenclature based on phylogenetic principles for SARS-CoV-2, but continuously assesses logistic growth rates, immune escape in comparison to BA.2, and mutational fitness per lineage^50^. Similarly, PyR0 is a hierarchical Bayesian multinomial logistic regression model that detects lineages increasing in prevalence as well as identifies mutations relative to lineage fitness^24^ and Episcore, predicts which existing amino acid mutations might contribute to future SARS-CoV-2 VOC’s^51^, though neither is run continuously nor available as a web-based platform. Other web-based platforms, such as CoVariants, provide an overview of variant frequencies and shared amino acid mutations^52^, and CovidCG tracks viral mutations, lineages, and clades in different countries over time^53^. CoVRadar focuses on mutation frequency by location and mutation distribution among sequences for the molecular surveillance of the SARS-CoV-2 spike protein^54^. Outbreak.info also provides information about lineages and amino acid changes while also reporting prevalence of variants, their geographical distributions and comparisons of changes between lineages^38^.

CoVerage remains unique in its application as it continuously monitors for variants with a potential selective advantage for further study. Only the EVEscape platform developed by Thadani and colleagues, similarly scores emerging variants by their immune escape potential^55^, however, does not directly identify variants with a potential selective advantage irrespective of this. It is this that makes CoVerage unique: its ability to identify variants with a potential selective advantage using lineage frequency dynamics in combination with predictions of lineage antigenic alterations. This allowed it to accurately identify 31 out of 35 (89%) VOCs and VOIs designated by the WHO with an average lead time of 84 days. We demonstrate the value of this approach in our comprehensive assessment of the value of such predictions for the early identification of past VOCs, VOIs and VUMs, with a lead time of almost three months relative to their maximal abundances and a precision of 79% and recall of 73%. As such, CoVerage lineage predictions are a valuable source of information for further investigations in clinical and epidemiological studies, allowing to prioritize lineages regarding their potential relevance for public health decision making and vaccine updates. The identification of amino acid changes throughout the spike protein that may have selective advantage can further inform antibody design and help understand the molecular basis of adaptive evolution^56^. Additionally, CoVerage links to alternative web-based resources for additional information on these selected lineages, providing a comprehensive resource for lineage surveillance. Each of the different analyses provided on CoVerage offers its own benefits when used individually and more so in combination with one another.

CoVerage provides a continuously updated resource for the *in silico* detection and characterization of potential VOIs, VOCs and VUMs from SARS-CoV-2 genomic surveillance data, to support researchers and public health officials in assessing and interpreting such data. In the future, the framework will be extended to provide relevant surveillance for other rapidly evolving viruses, such as seasonal influenza viruses, and to integrate further data types, such as metagenomic sequences of wastewater samples^57^.

## Methods

### Lineage dynamics by region

In the sublineage corrected analysis, the lineage frequencies are corrected by including isolates belonging to sublineages in the count, and subsequently processed as before. For example, three sublineages BA.1.2, BA.1.3 and BA.1.4 will be summarized into BA.1 and then analyzed with Fisher’s exact test. This approach leads to a better resolution of the lineage branches likely associated with more rapid spread and lineages with a selective advantage split into several sublineages can be better detected. Here Pango lineages were used as nomenclature as they define an epidemiologically relevant phylogenetic cluster in which new lineages are only designated if the lineage has high coverage and contains a sufficient number of sequences^21^.

Lastly, a sliding window approach was implemented to achieve a more sensitive detection than the monthly analysis. For this analysis, sequences are sorted by date and frequencies in the *w* sequences in the current window are compared to the *w* previous ones using Fisher’s exact test. The window is moved over the data using a step size *s*. Significance estimates are corrected for multiple testing through correction of p-values of the multiple tests with Benjamini-Yekutieli procedure^58^. For the analysis on countries, *w* = 1000 and *s* = 100 are used and for the analysis on more granular German state level *w* = 200 and *s* = 10 is chosen. Additionally, we require a significant (FDR < 0.05) increase for a certain window and a frequency threshold of 0.1 to report results. The reported date, or the date of the significant increase, is the date of the last sequence in the current window.

### Spike protein allele dynamics

To identify amino acid changes in the spike protein that may provide a lineage with a selective advantage, the sweep dynamic (SD) plot method^17,20^ is used on spike protein sequences extracted from viral genome sequences data downloaded from GISAID and Zenodo. To execute this methodology on the large SARS-CoV-2 sequence collection, 500 sequences per month per country are subsampled randomly and downloaded with GISAIDR^59^, and subsequently, identical sequences per time period are clustered using CD-HIT^60^. For German state-wise analysis, we randomly downloaded 2000 sequences per month for the entire country and divided them by states. Next, a multiple sequence alignment is generated using MAFFT^61^, with the spike sequence of Wuhan/IPBCAMS-WH-01/2019 as a reference. Phylogenetic trees are inferred for each country using fasttree^62^, and the Sankoff algorithm is applied for ancestral character state reconstruction, and as in Steinbrueck & McHardy, 2011, we use the same cost for all amino acid changes and indels^20^. We previously showed that in ancestral sequence reconstruction for the major antigen of human influenza A (H3N2) viruses, both approaches produce very similar results, due to the very small evolutionary time scales being considered^20^. The calculation of relative frequencies of specific spike protein alleles and statistical evaluation are then performed as described in Steinbrück & McHardy, 2011, with a frequency threshold of 0.1 for finding spike protein alleles and their amino acid changes that may provide a selective advantage^20^.

### *In silico* assessment of antigenic lineage alterations

The antigenic alteration score for a lineage *a*_*l*_ is calculated using a method we developed to score each SARS-CoV-2 spike protein sequence relative to the Wuhan-Hu-1 strain (Additional File 1). Antigenic weights were applied to the amino acid changes occurring throughout the spike protein based on a viral immune escape scoring matrix derived from an antigenic tree for seasonal Influenza A (H3N2) viruses (IAV)^56^. The weights were then summed per isolate and then averaged across all isolates of a given circulating Pango Lineage to calculate the final antigenic score, *a*_*l*_. To select lineages considered significantly altered antigenically for a given month, a z-score was applied to each circulating Pango lineage for the given month with a frequency of 0.001 or greater. To calculate the z-score, the population mean, or the mean of the lineages circulating that month above the frequency threshold was subtracted from the individual lineage score and then divided by the standard deviation of those lineages circulating above the frequency threshold. Circulating lineages with an assigned z-score of one or greater were considered to be significantly antigenically altered. The threshold of 1 was chosen to distinguish VOCs and VOIs designated until June 2024 from other lineages (**Fig. 5c**).

In addition to the individual lineage score, a country-wide antigenic alteration score *a*_*c*_ is calculated by summing over all antigenic lineage scores, weighted by their frequencies for a particular period in time:

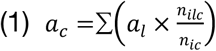

Here, *n*_*ilc*_ is the number of isolates of each Pango lineage for country *c* and *n*_*ic*_ is the total number of isolates per country.

## Data Availability

The viral genome sequences and metadata for conducting *in silico* assessment of antigenic lineage alterations that were used to generate the heatmaps and global antigenic maps for the months of November 2021 through January 2022 and January 2023 to March 2023 are publicly available from GISAID (http://gisaid.org). GISAID IDs are available at https://github.com/hzi-bifo/corona_lineage_dynamics/data/ and the results can be found on Zenodo (https://zenodo.org/records/10171227).

## Code Availability

All the codes are publicly available at: https://github.com/hzi-bifo/corona_lineage_dynamics, https://github.com/hzi-bifo/Corona_Variant_Scoring and https://github.com/hzi-bifo/corona_protein_dynamics.

## Acknowledgements

We gratefully acknowledge all data contributors, i.e. the Authors and their Originating laboratories responsible for obtaining the specimens, and their Submitting laboratories for generating the genetic sequence and metadata and sharing via the GISAID Initiative, on which this research is based. A.C.M. gratefully acknowledges funding from the German Centre for Infection Research project TI 12.002.

## Author contributions

K.N., Z-L.D., G.R., S.G., M.H.F.A., M.H., and S.R. developed the methods and wrote the code. K.N., S.R., and A.C.M. analysed and interpreted the data, K.N., S.R., and A.C.M. wrote the paper and A.C.M. designed the research study. M.H. suggested data visualizations and commented on the manuscript, K.N., Z-L.D., G.R., M.H.F.A., and S.G. maintain the web service. All authors read and approved the manuscript.

