## Supplementary Information for "*In silico* genomic surveillance by CoVerage predicts and characterizes SARS-CoV-2 Variants of Interest"

Provided in this supplementary material is an example of the CoVerage homepage (**Supplementary Fig. 1**) and an allele dynamics plot for the USA (**Supplementary Figure 2**) along with reference material used in the variant antigenic alteration scoring analysis (**Supplementary Table 1**) and the resultant antigenic alteration scores for the month of March 2023 (**Supplementary Table 2**). The circulating variants for the month of March 2023 are given with their associated antigenic alteration scores and ranked from highest to lowest for that month. Also included are the results for the lineage dynamics analysis for the months of January 2023 through March 2023 (**Supplementary Tables 3-5**). These tables list the selected pVOIs and their median p-value along with their WHO designation as either a variant under monitoring (VUM) or a variant of concern (VOC).

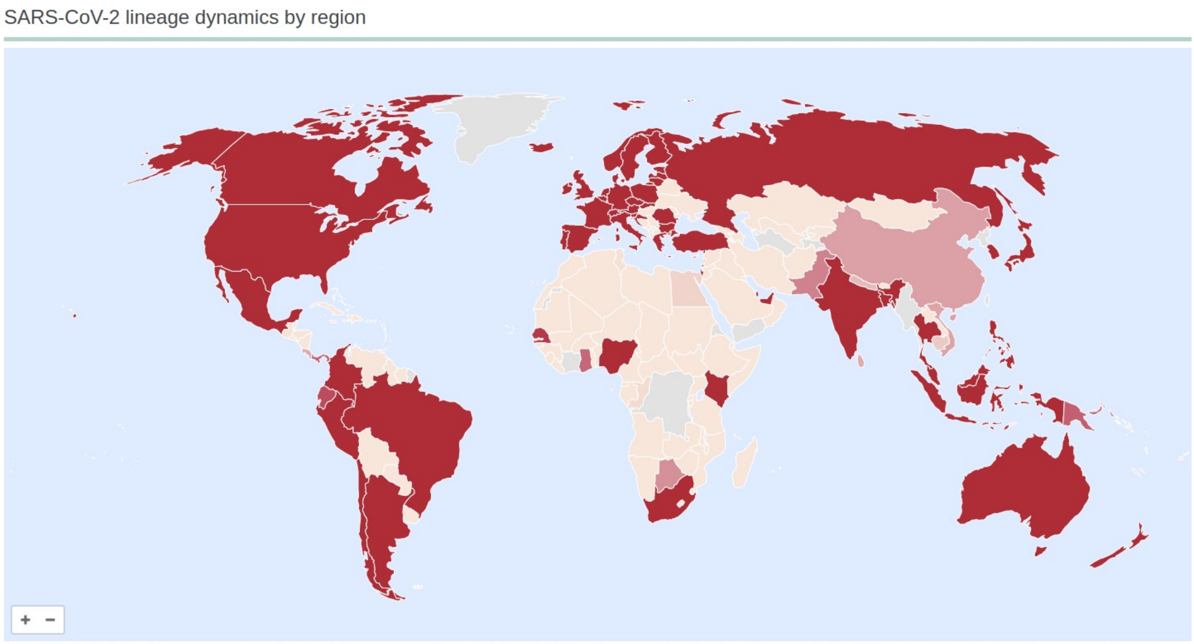


**Supplementary Fig. 1: Global map representing the SARS-CoV-2 lineage dynamics by region on the CoVerage homepage.** Countries shown in red have more than 2000 sequences and have the lineage dynamics analysis results available. Case numbers per country are obtained from the WHO Coronavirus (COVID-19) data repository[20].

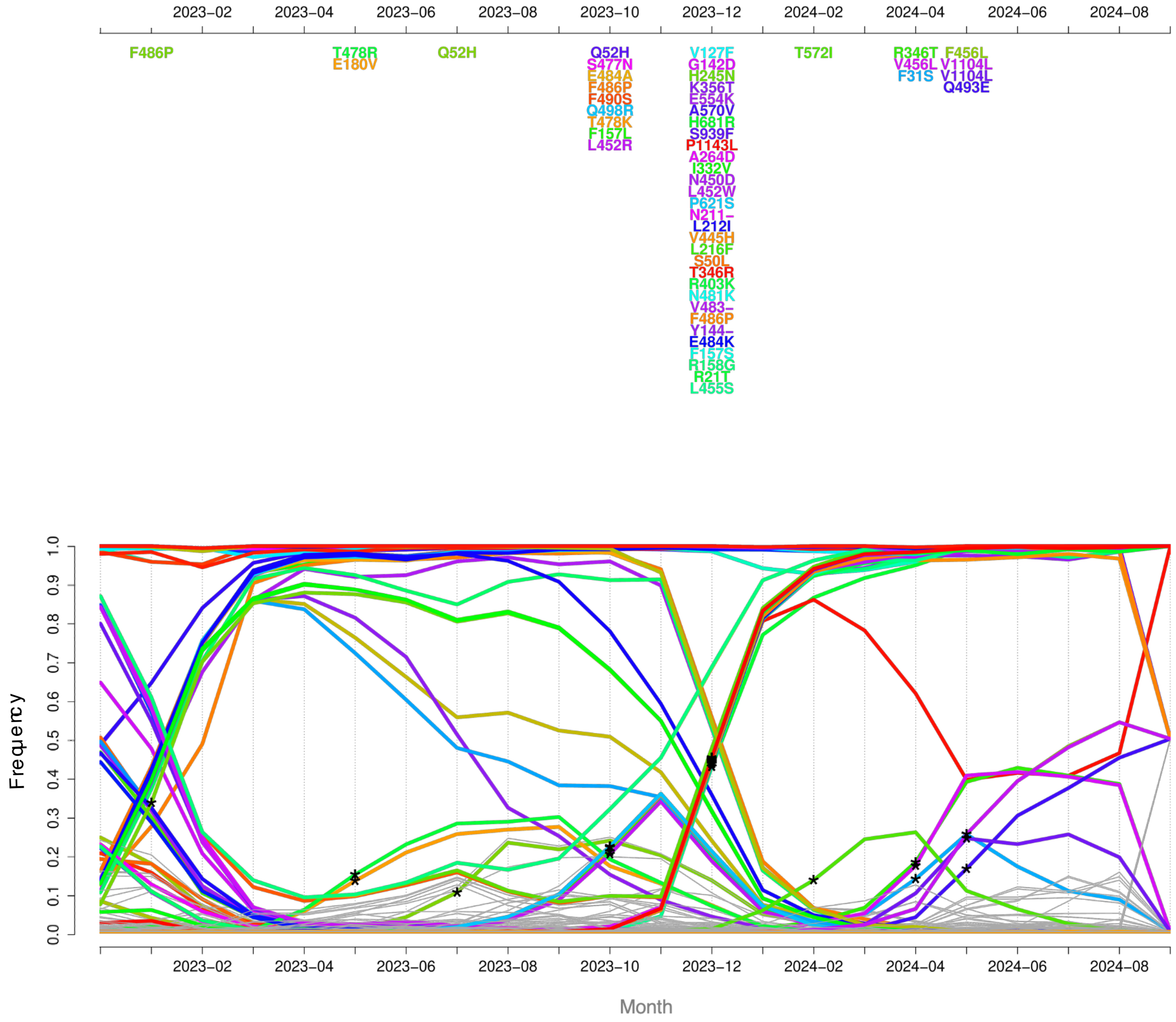


**Supplementary Fig. 2: b SD plot for United States of America from December 2022 to July 2024.** The asterisks represent when a spike protein allele with certain amino acid changes significantly rises in frequency and the color of the curve corresponds to the associated amino acid change. Amino acid changes are specified at the top of the plot, based on the time point when they were identified as significant.

**Supplementary Table 1: Selected amino acid sites that have been shown to alter SARS-CoV-2 antigenicity using literature available up to April 2021.** Antigenic sites are located on the SARS-CoV-2 spike protein and have demonstrated a capacity to alter the antigenicity of the virus.

| antigenic_sites | information |
|---|---|
| 18 | escape from NTD-binding mAbs, reduces antibody neutralization |
| 157 | reduces neutralization by mAb (2489) |
| 234 | resistant to neutralizing antibodies |
| 417 | escape neutralization by mAbs, enhanced binding with ACE2 |
| 439 | increased binding affinity for ACE2 through formation of new salt bridge, resistant to some neutralizing antibodies |
| 440 | immune escape the neutralization by mAbs |
| 444 | immune escape the neutralization by mAbs and human convalescent sera |
| 446 | immune escape the neutralization by mAbs and human convalescent sera |
| 450 | immune escape the neutralization by mAbs and human convalescent sera |
| 452 | reduces neutralization by mAbs, enhances infectivity |
| 453 | reduces neutralization by mAbs, increase ACE2 affinity |
| 475 | immune escape the neutralization by mAbs and human convalescent sera |
| 476 | immune escape the neutralization by mAbs |
| 477 | immune escape, resistant to neutralization, enhanced binding with ACE2 |
| 478 | immune escape the neutralization by mAbs and human convalescent sera |
| 479 | immune escape the neutralization by mAbs |
| 483 | resistant to some neutralizing antibodies and to human convalescent sera |
| 484 | reduces neutralization of antibodies, immune escape |
| 486 | immune escape the neutralization by mAbs |
| 489 | immune escape the neutralization by mAbs |
| 490 | resistant to some neutralizing antibodies |
| 493 | immune escape the neutralization by mAbs |
| 499 | immune escape the neutralization by mAbs and human convalescent sera |
| 501 | reduces neutralization of RBD antibodies, increases ACE2 binding affinity, increases transmissibility |
| 681 | immune escape the neutralization by mAbs and human convalescent sera |
| 769 | decreased susceptibility to neutralizing antibodies |
| 796 | reduction in susceptibility to non-RBD specific antibodies, but decreases infectivity |

**Supplementary Table 2: Antigenic alteration scores for circulating lineages in the month of March 2023.** Given are the Pango lineages, their associated antigenic alteration score, and their rank among circulating lineages of that month.

| Pango lineage | Antigenic score | rank | Pango lineage | Antigenic score | rank | Pango lineage | Antigenic score | rank |
|---|---|---|---|---|---|---|---|---|
| CM.8 | 6.979 | 1 | DJ.1 | 5.671 | 117 | BN.1.3 | 5.184 | 233 |
| CM.2.1 | 6.950 | 2 | BQ.1.13 | 5.664 | 118 | BA.5.2.34 | 5.181 | 234 |
| CM.11 | 6.946 | 3 | BA.5.2.18 | 5.655 | 119 | XBC.2 | 5.175 | 235 |
| CM.4 | 6.493 | 4 | BQ.1.1.28 | 5.643 | 120 | CV.1 | 5.170 | 236 |
| CM.7 | 6.395 | 5 | DJ.1.1 | 5.642 | 121 | BE.8 | 5.170 | 237 |
| CM.8.1 | 6.347 | 6 | BA.5.1.23 | 5.639 | 122 | XBK | 5.166 | 238 |
| CB.1 | 6.326 | 7 | XAY.2 | 5.623 | 123 | BA.1.1 | 5.163 | 239 |
| BF.33 | 6.325 | 8 | BE.1.1.1 | 5.618 | 124 | BN.1.2.1 | 5.157 | 240 |
| BA.4.6.3 | 6.325 | 10 | BQ.1.23 | 5.612 | 125 | BE.1.2.1 | 5.156 | 241 |
| BA.5.1.22 | 6.325 | 10 | BQ.1.10.1 | 5.605 | 126 | BN.1.4 | 5.154 | 242 |
| CK.3 | 6.325 | 10 | BQ.1.8 | 5.603 | 127 | XBF | 5.151 | 243 |
| CM.12 | 6.254 | 13.5 | BA.5.2.1 | 5.600 | 128 | BF.7.5 | 5.147 | 244 |
| CM.1 | 6.254 | 13.5 | BQ.1.28 | 5.599 | 129 | BA.5.1 | 5.140 | 245 |
| XBJ | 6.254 | 13.5 | BA.5.2.33 | 5.592 | 130 | BL.6 | 5.139 | 246 |
| CM.5 | 6.254 | 13.5 | BQ.1.9 | 5.580 | 131 | BN.1.7 | 5.139 | 247 |
| BM.2.1 | 6.216 | 16 | BQ.1.25 | 5.561 | 132 | BA.4.6 | 5.137 | 248 |
| CM.2 | 6.198 | 17 | BQ.1.18 | 5.556 | 133 | BF.11 | 5.118 | 249 |
| CK.1.1 | 6.143 | 18 | BQ.1.1.11 | 5.553 | 134 | BY.1 | 5.116 | 250 |
| BA.5.2.19 | 6.142 | 19.5 | BF.7.8 | 5.547 | 135 | BN.1.3.1 | 5.106 | 251 |
| BA.5.2.43 | 6.142 | 19.5 | BQ.1.1.10 | 5.543 | 136 | BQ.1.17 | 5.096 | 252 |
| CK.2.1 | 6.123 | 21 | BQ.1.1.21 | 5.532 | 137 | BA.5.2 | 5.095 | 253 |
| BQ.1.26 | 6.108 | 22 | BQ.1 | 5.531 | 138 | BN.1.3.3 | 5.066 | 254 |
| CK.1 | 6.106 | 23 | XBB.6 | 5.528 | 139 | DF.1.1 | 5.065 | 255.5 |
| BQ.1.16 | 6.081 | 24 | CQ.1 | 5.524 | 140 | BF.7.9 | 5.065 | 255.5 |
| BA.2.3.20 | 6.070 | 25 | BA.5.2.12 | 5.523 | 141 | BF.7.3 | 5.057 | 257 |
| DR.1 | 6.063 | 26 | BQ.1.1.19 | 5.509 | 142 | BE.1.1 | 5.028 | 258 |
| DT.1 | 6.015 | 33.5 | CN.1 | 5.499 | 143 | BA.1.17 | 5.007 | 259 |
| BQ.1.20 | 6.015 | 33.5 | CH.1.1 | 5.497 | 144 | BA.2.10.4 | 5.003 | 260.5 |
| BQ.1.1.26 | 6.015 | 33.5 | CQ.2 | 5.484 | 145 | BA.2.12 | 5.003 | 260.5 |
| BQ.1.8.1 | 6.015 | 33.5 | XBC.1.2.1 | 5.484 | 146 | Unassigned | 4.999 | 262 |
| BA.5.2.25 | 6.015 | 33.5 | BA.5.1.5 | 5.464 | 147 | BA.2.82 | 4.997 | 263.5 |
| BA.5.2.46 | 6.015 | 33.5 | XAY.1.1 | 5.459 | 148 | BA.2.14 | 4.997 | 263.5 |
| BQ.1.6 | 6.015 | 33.5 | BA.5.2.13 | 5.435 | 149 | BF.25 | 4.981 | 265 |
| BA.5.6.2 | 6.015 | 33.5 | CR.1 | 5.422 | 150 | BA.4.1 | 4.970 | 266 |
| DE.2 | 6.015 | 33.5 | BF.7.4 | 5.402 | 151 | BA.5.1.26 | 4.959 | 267 |
| DM.1 | 6.015 | 33.5 | BA.2.75.4 | 5.394 | 152 | BQ.1.19 | 4.952 | 268 |
| BQ.1.1.14 | 6.015 | 33.5 | XBC.1.1 | 5.393 | 153 | BN.1.1 | 4.951 | 269 |
| BQ.1.8.2 | 6.015 | 33.5 | CH.1 | 5.371 | 154 | BA.2.10 | 4.938 | 270 |
| BQ.1.1.9 | 6.015 | 33.5 | BQ.1.27 | 5.369 | 155 | CM.10 | 4.935 | 271 |
| BE.1.2 | 6.015 | 33.5 | BU.1 | 5.354 | 156 | BQ.1.25.1 | 4.926 | 272 |
| BM.1.1.5 | 6.012 | 41 | BA.2.75.5 | 5.352 | 157 | BA.3.1 | 4.888 | 273 |
| CH.3.1 | 6.012 | 42 | BA.5 | 5.345 | 158 | CA.3.1 | 4.882 | 274 |
| BQ.1.1.30 | 6.006 | 43 | BA.5.2.48 | 5.332 | 159 | BM.4.1.1 | 4.870 | 275 |
| BQ.1.22 | 5.998 | 44 | XBB.1.1 | 5.326 | 160 | XBB.1.2 | 4.854 | 276 |

| | | | | | | | | |
|---|---|---|---|---|---|---|---|---|
| DN.1.1 | 5.997 | 45 | BQ.1.4 | 5.321 | 161 | XBL | 4.853 | 277 |
| BQ.1.13.1 | 5.992 | 46 | BA.5.3.3 | 5.317 | 179 | BA.5.3.5 | 4.833 | 278 |
| BQ.1.1.34 | 5.982 | 47 | BF.28 | 5.317 | 179 | XBC.1 | 4.822 | 279 |
| BY.1.1.1 | 5.979 | 48 | DF.1 | 5.317 | 179 | BN.1.5 | 4.820 | 280 |
| DU.1 | 5.968 | 49 | BA.4.6.1 | 5.317 | 179 | BE.5 | 4.802 | 281 |
| CH.1.1.3 | 5.967 | 50 | XBG | 5.317 | 179 | CR.2 | 4.768 | 282 |
| CL.1 | 5.957 | 51 | BF.11.3 | 5.317 | 179 | BN.2 | 4.758 | 283 |
| BQ.1.1.32 | 5.952 | 52 | BF.11.5 | 5.317 | 179 | XBB.6.1 | 4.751 | 284 |
| BQ.1.21 | 5.941 | 53 | BF.5.1 | 5.317 | 179 | XBB.1.4 | 4.747 | 285 |
| CH.1.1.4 | 5.940 | 54 | BF.11.1 | 5.317 | 179 | XBB.1.5 | 4.724 | 286 |
| BQ.1.26.1 | 5.931 | 55 | BA.5.2.7 | 5.317 | 179 | BR.1 | 4.724 | 287 |
| CK.2.1.1 | 5.922 | 56 | BA.5.1.17 | 5.317 | 179 | BR.2.1 | 4.720 | 288 |
| BA.5.5.1 | 5.920 | 58.5 | CN.2 | 5.317 | 179 | XBB.1.9.1 | 4.712 | 289 |
| BA.5.2.37 | 5.920 | 58.5 | BA.5.2.39 | 5.317 | 179 | BR.2 | 4.707 | 290 |
| BA.5.2.32 | 5.920 | 58.5 | BA.5.1.6 | 5.317 | 179 | BA.1.18 | 4.706 | 291 |
| BU.3 | 5.920 | 58.5 | CP.1 | 5.317 | 179 | BM.1.1.1 | 4.698 | 292 |
| BQ.1.1.24 | 5.914 | 61 | BF.29 | 5.317 | 179 | BW.1.1 | 4.663 | 293 |
| BA.2.56 | 5.912 | 62 | BF.7.10 | 5.317 | 179 | BN.1.4.1 | 4.643 | 294 |
| BA.5.2.27 | 5.904 | 63 | BA.5.1.20 | 5.317 | 179 | BA.2 | 4.632 | 295 |
| BQ.1.1.31 | 5.901 | 64 | BE.7 | 5.317 | 179 | XBB.1.4.1 | 4.564 | 296 |
| BQ.1.11 | 5.889 | 65 | BF.11.2 | 5.317 | 179 | B.1.177 | 4.563 | 297 |
| BQ.1.1.16 | 5.883 | 66 | BA.5.1.24 | 5.317 | 179 | CP.2 | 4.558 | 298 |
| BQ.1.1.6 | 5.880 | 67 | BA.5.1.30 | 5.317 | 179 | XBB.1.9 | 4.553 | 299 |
| BQ.1.15 | 5.877 | 68 | BA.5.2.26 | 5.317 | 179 | XBB.1.8 | 4.534 | 300 |
| BQ.1.1.22 | 5.868 | 69 | BF.7.13.2 | 5.317 | 179 | CA.1 | 4.529 | 301 |
| BA.2.12.1 | 5.856 | 70 | BA.5.2.35 | 5.317 | 179 | XBB.2 | 4.527 | 302 |
| BQ.1.12 | 5.854 | 71 | BA.5.2.3 | 5.317 | 179 | XBB.3 | 4.515 | 303 |
| BQ.1.1.29 | 5.846 | 72 | CR.1.3 | 5.317 | 179 | BS.1.1 | 4.478 | 304 |
| BQ.1.24 | 5.845 | 73 | BA.5.1.18 | 5.317 | 179 | BA.1 | 4.412 | 305 |
| CG.1 | 5.842 | 74 | BF.2 | 5.317 | 179 | XBB.2.2 | 4.352 | 306 |
| XBB.2.1 | 5.839 | 75 | BF.7.1 | 5.317 | 179 | BA.5.1.12 | 4.351 | 307 |
| BQ.1.1.25 | 5.834 | 76 | BE.6 | 5.317 | 179 | BA.5.2.24 | 4.319 | 308 |
| DN.1 | 5.828 | 77 | BF.7.5.1 | 5.317 | 179 | BA.5.2.14 | 4.310 | 309 |
| BQ.1.1.27 | 5.827 | 78 | CP.3 | 5.317 | 179 | BR.1.2 | 4.307 | 310 |
| CH.1.1.5 | 5.825 | 79 | BA.5.9 | 5.317 | 179 | B.1.617.2 | 4.289 | 311 |
| BQ.1.1.2 | 5.823 | 80 | BA.5.1.28 | 5.317 | 179 | BR.3 | 4.286 | 312 |
| DG.1 | 5.821 | 81.5 | BF.7.4.1 | 5.316 | 197 | BR.3 | 4.286 | 312 |
| CK.1.2 | 5.821 | 81.5 | BF.5 | 5.316 | 198 | BA.2.61 | 4.277 | 313.5 |
| BF.14 | 5.820 | 83 | BF.7.14 | 5.316 | 199 | BE.1.1.2 | 4.277 | 313.5 |
| DS.1 | 5.816 | 84 | BN.1.6 | 5.314 | 200.5 | BA.2.3 | 4.195 | 315 |
| BW.1 | 5.813 | 85 | BN.1.8 | 5.314 | 200.5 | BJ.1 | 4.160 | 316 |
| BE.9 | 5.813 | 86 | BF.7.15 | 5.312 | 202 | XBB.1 | 4.156 | 317 |
| BQ.1.1.1 | 5.812 | 87 | BN.1.3.2 | 5.311 | 203 | BA.2.75 | 4.151 | 318 |
| BQ.1.1.15 | 5.812 | 88 | BA.5.2.49 | 5.301 | 204 | BM.2 | 4.075 | 319 |
| CH.1.1.1 | 5.811 | 89 | XBB.4 | 5.294 | 205 | BN.1.9 | 4.045 | 320 |
| BM.1.1.3 | 5.809 | 90 | BA.5.2.6 | 5.290 | 206 | BA.2.21 | 4.037 | 321 |
| BQ.1.1.7 | 5.800 | 91 | CJ.1 | 5.287 | 207 | BA.5.2.28 | 4.027 | 322 |
| BQ.1.1.8 | 5.786 | 92 | CH.3 | 5.286 | 208 | BA.2.1 | 3.990 | 323 |
| DB.1 | 5.782 | 93 | BE.1.4 | 5.281 | 209 | XBB | 3.943 | 324 |
| BQ.1.1.3 | 5.778 | 94 | CR.1.1 | 5.272 | 210 | BA.5.2.16 | 3.886 | 325 |

| | | | | | | | | |
|---|---|---|---|---|---|---|---|---|
| BE.4 | 5.776 | 96 | BA.2.9 | 5.260 | 211 | BF.26 | 3.884 | 326 |
| BE.4.1.1 | 5.776 | 96 | BF.7.6 | 5.259 | 212 | XBB.1.7 | 3.832 | 327 |
| BF.16 | 5.776 | 96 | CM.5.2 | 5.246 | 213 | BM.1 | 3.812 | 328 |
| BQ.1.1.20 | 5.771 | 98 | BE.10 | 5.237 | 214 | BA.5.1.10 | 3.811 | 329 |
| BQ.1.1.23 | 5.771 | 99 | BN.1.1.1 | 5.236 | 215 | BA.5.1.3 | 3.605 | 330 |
| BQ.1.1.13 | 5.768 | 100 | BF.7 | 5.234 | 216 | BM.1.1 | 3.538 | 331 |
| BN.3.1 | 5.763 | 101 | BN.1 | 5.230 | 217 | BE.1 | 3.512 | 332 |
| BQ.1.1.5 | 5.761 | 102 | BM.1.1.4 | 5.229 | 218 | BM.4 | 3.416 | 333 |
| BQ.1.1.18 | 5.754 | 103 | BE.4.2 | 5.220 | 219 | BA.5.2.21 | 3.252 | 334 |
| BQ.1.2 | 5.746 | 104 | CJ.1.1 | 5.210 | 220 | BA.1.1.7 | 3.231 | 335 |
| BM.4.1 | 5.745 | 105 | BA.5.1.27 | 5.205 | 221 | BA.2.10.1 | 2.912 | 336 |
| CM.4.1 | 5.740 | 106 | BN.1.2 | 5.203 | 222 | BA.3 | 1.530 | 337 |
| BQ.1.1.17 | 5.731 | 107 | CA.7 | 5.202 | 223 | BA.2.73 | 1.448 | 339 |
| BA.5.2.20 | 5.730 | 108 | BA.5.11 | 5.200 | 224 | DP.1 | 1.448 | 339 |
| BQ.1.1.4 | 5.726 | 109 | BA.1.17.2 | 5.198 | 226.5 | XAS | 1.448 | 339 |
| BQ.1.10 | 5.722 | 110 | BA.1.13 | 5.198 | 226.5 | B.1.1 | 0.694 | 341 |
| BQ.1.14 | 5.720 | 111 | BA.1.15.1 | 5.198 | 226.5 | AY.36 | 0.513 | 343 |
| BQ.1.3 | 5.716 | 112 | BA.1.21.1 | 5.198 | 226.5 | AY.122 | 0.513 | 343 |
| CH.1.1.2 | 5.699 | 113 | XAY.1 | 5.195 | 229 | AY.43 | 0.513 | 343 |
| BQ.1.5 | 5.691 | 114 | BA.1.1.18 | 5.192 | 230 | B | 0.000 | 345.5 |
| BQ.1.1 | 5.686 | 115 | BA.5.2.47 | 5.189 | 231 | B.1 | 0.000 | 345.5 |
| BA.5.3.1 | 5.673 | 116 | BF.13 | 5.187 | 232 | | | |

**Supplementary Table 3: Selected pVOIs listed by their Pango lineage for January 2023 and their median p-values as per the lineage dynamics analysis.** Listed p-values are the result of the Fisher's exact test to determine lineages significantly on the rise in frequency. Also shown is the data and designation of lineages as a VOI or Variant Under Monitoring (VUM).

| Pango lineage | Median p-value | Identified as a VOC / VOI / VUM | Notes (dates given as DD-MM-YYYY) | References |
|---|---|---|---|---|
| BA.1.1 | 9.60E-63 | | | |
| BA.2 | 4.94E-55 | | de-escalated VOC as per the ECDC | https://www.ecdc.europa.eu/en/covid-19/variants-concern |
| BA.2.10.1 | 0.005404522 | | | |
| BA.2.3 | 9.27E-26 | | | |
| BA.2.3.20 | 6.39E-67 | | | |
| BA.4.6 | 9.11E-11 | | enhanced neutralization resistance respective of parental BA.4/5 subvariant | Qu. P et al., 2023 |
| BA.5 | 2.21E-28 | | de-escalated VOC as per the ECDC, broadly resistant to most nAbs | Cao et al., 2022; https://www.ecdc.europa.eu/en/covid-19/variants-concern |
| BA.5.1 | 7.16E-63 | | | |
| BA.5.11 | 7.41E-05 | | | |
| BA.5.2 | 6.44E-15 | | | |
| BA.5.2.1 | 3.25E-10 | | | |
| BA.5.2.6 | 1.48E-09 | | | |
| BE.1.1 | 3.08E-66 | | | |
| BE.9 | 1.60E-65 | | | |
| BF.5 | 7.59E-280 | | | |
| BF.7 | 2.41E-12 | | enhanced neutralization resistance respective of parental BA.4/5 subvariant | Qu. P et al., 2023 |
| BF.7.14 | 6.84E-31 | | | |
| BN.1.2 | 1.97E-15 | | | |
| BN.1.3 | 2.69E-16 | | | |
| BQ.1 | 5.46E-14 | VOI (20-10-2022) | VOI as per ECDC (20-10-2022); enhanced neutralization resistance respective of parental BA.5 subvariant | Qu. P et al., 2023; https://www.ecdc.europa.eu/en/publications-data/spread-sars-cov-2-omicron-variant-sub-lineage-bq1-eueea |
| BQ.1.1 | 3.25E-25 | | resistant to all clinical mAbs / enhanced neutralization resistance respective of parental BA.5 subvariant | Arora P., et al. 2022; Qu P., et al., 2023 |
| BQ.1.1.1 | 3.06E-08 | | | |
| BQ.1.1.10 | 0.003182437 | | | |
| BQ.1.1.20 | 6.32E-82 | | | |
| BQ.1.1.22 | 3.55E-06 | | | |
| BQ.1.1.38 | 0.000723485 | | | |
| BQ.1.1.4 | 0.031009802 | | | |
| BQ.1.10 | 1.80E-05 | | | |
| BR.2.1 | 6.67E-159 | | | |

| | | | | |
|---|---|---|---|---|
| CH.1.1 | 0.026651959 | VUM (08-02-2023) | WHO variant under monitoring since (08-02-2023) | https://www.who.int/activities/tracking-SARS-CoV-2-variants |
| CH.1.1.1 | 0.000159952 | | | |
| CH.1.1.7 | 7.98E-18 | | | |
| CK.1 | 1.18E-05 | | | |
| CL.1 | 5.48E-95 | | | |
| CM.12 | 2.29E-07 | | | |
| DY.2 | 5.38E-08 | | | |
| DY.4 | 5.12E-15 | | | |
| Unassigned | 0.029477699 | | | |
| XBB | 1.42E-58 | VUM (12-10-2022) | XBB and its sublineages have been identified as variants under monitoring as per the WHO (12-10-2022) | https://www.who.int/activities/tracking-SARS-CoV-2-variants |
| XBB.1 | 1.60E-05 | | | |
| XBB.1.15 | 0.002277383 | | | |
| XBB.1.5 | 4.52E-22 | VOI (11-01-2023) | WHO current variant of interest (as of 11-01-2023) | https://www.who.int/activities/tracking-SARS-CoV-2-variants |
| XBB.1.5.12 | 2.84E-05 | | | |
| XBB.1.9.1 | 3.02E-05 | VUM (30-03-2023) | WHO variant under monitoring since (30-03-2023) | https://www.who.int/activities/tracking-SARS-CoV-2-variants |
| XBB.1.9.2 | 2.18E-07 | VUM (26-04-2023 | WHO variant under monitoring since (26-04-2023) | https://www.who.int/activities/tracking-SARS-CoV-2-variants |
| XBB.2 | 8.09E-58 | | | |
| XBB.2.6 | 1.24E-115 | | | |
| XBF | 2.68E-08 | | increased antibody neutralization | Ackerman A., et al. 2023 |

**Supplementary Table 4: Selected pVOIs listed by their Pango lineage for February 2023 and their median p-values as per the lineage dynamics analysis.** Listed p-values are the result of the Fisher's exact test to determine lineages significantly on the rise in frequency. Also shown is the data and designation of lineages as a VOI or Variant Under Monitoring (VUM).

| Pango lineage | Median p-value | Identified as a VOC / VOI / VUM | Notes (dates given as DD-MM-YYYY) | References |
|---|---|---|---|---|
| BA.2 | 4.94E-55 | | de-escalated VOC as per the ECDC | https://www.ecdc.europa.eu/en/covid-19/variants-concern |
| BA.2.10.1 | 0.005404522 | | | |
| BA.2.75 | 1.78E-200 | VUM (06-07-2022) | WHO variant under monitoring since (06-07-2022), VOI as per the ECDC | https://www.who.int/activities/tracking-SARS-CoV-2-variants; https://www.ecdc.europa.eu/en/covid-19/variants-concern |
| BA.4.1.9 | 0.012054399 | | | |
| BA.5.2 | 6.44E-15 | | | |
| BA.5.2.1 | 3.25E-10 | | | |
| BA.5.2.6 | 1.48E-09 | | | |
| BE.9 | 1.60E-65 | | | |
| BF.7 | 2.41E-12 | | enhanced neutralization resistance respective of parental BA.4/5 subvariant | Qu. P et al., 2023 |
| BF.7.14 | 6.84E-31 | | | |
| BN.1.2 | 1.97E-15 | | | |
| BN.1.3 | 2.69E-16 | | | |
| BQ.1 | 5.46E-14 | VOI (20-10-2022) | VOI as per ECDC (20-10-2022); enhanced neutralization resistance respective of parental BA.5 subvariant | Qu. P et al., 2023; https://www.ecdc.europa.eu/en/publications-data/spread-sars-cov-2-omicron-variant-sub-lineage-bq1-eueea |
| BQ.1.1 | 3.25E-25 | | resistant to all clinical mAbs / enhanced neutralization resistance respective of parental BA.5 subvariant | Arora P., et al. 2022; Qu P., et al., 2023 |
| BQ.1.1.1 | 3.06E-08 | | | |
| BQ.1.1.10 | 0.003182437 | | | |
| BQ.1.1.4 | 0.031009802 | | | |
| BQ.1.10 | 1.80E-05 | | | |
| BR.2.1 | 6.67E-159 | | | |
| CH.1.1 | 0.026651959 | VUM (08-02-2023) | WHO variant under monitoring since (08-02-2023) | https://www.who.int/activities/tracking-SARS-CoV-2-variants |
| CH.1.1.1 | 0.000159952 | | | |
| CH.1.1.7 | 7.98E-18 | | | |
| CK.1 | 1.18E-05 | | | |
| CL.1 | 5.48E-95 | | | |
| CM.12 | 2.29E-07 | | | |
| DY.2 | 5.38E-08 | | | |
| DY.4 | 5.12E-15 | | | |
| EG.1 | 2.85E-08 | | | |

| | | | | |
|---|---|---|---|---|
| FL.2 | 1.08E-22 | | | |
| FR.1 | 0.020160177 | | | |
| Unassigned | 0.029477699 | | | |
| XBB | 1.42E-58 | VUM (12-10-2022) | XBB and its sublineages have been identified as variants under monitoring as per the WHO (12-10-2022) | https://www.who.int/activities/tracking-SARS-CoV-2-variants |
| XBB.1 | 1.60E-05 | | | |
| XBB.1.15 | 0.002277383 | VOI (11-01-2023) | WHO current variant of interest (as of 11-01-2023) | https://www.who.int/activities/tracking-SARS-CoV-2-variants |
| XBB.1.16 | 5.97E-10 | VOI (17-04-2023) | WHO current variant of interest (as of 17-04-2023) | https://www.who.int/activities/tracking-SARS-CoV-2-variants |
| XBB.1.16.1 | 0.006501881 | | | |
| XBB.1.18.1 | 2.35E-22 | | | |
| XBB.1.5 | 4.52E-22 | | | |
| XBB.1.5.12 | 2.84E-05 | | | |
| XBB.1.5.25 | 3.96E-41 | | | |
| XBB.1.5.33 | 5.67E-05 | | | |
| XBB.1.9.1 | 3.02E-05 | VUM (30-03-2023) | WHO variant under monitoring since (30-03-2023) | https://www.who.int/activities/tracking-SARS-CoV-2-variants |
| XBB.1.9.2 | 2.18E-07 | VUM (26-04-2023) | WHO variant under monitoring since (26-04-2023) | https://www.who.int/activities/tracking-SARS-CoV-2-variants |
| XBB.2 | 8.09E-58 | | | |
| XBB.2.6 | 1.24E-115 | | | |
| XBF | 2.68E-08 | | increased antibody neutralization | Ackerman A., et al. 2023 |

**Supplementary Table 5: Selected pVOIs listed by their Pango lineage for March 2023 and their median p-values as per the lineage dynamics analysis.** Listed p-values are the result of the Fisher's exact test to determine lineages significantly on the rise in frequency. Also shown is the data and designation of lineages as a VOI or Variant Under Monitoring (VUM).

| Pango lineage | Median p-value | Identified as a VOC / VOI / VUM | Notes (dates given as DD-MM-YYYY) | References |
|---|---|---|---|---|
| AY.43 | 7.02E-26 | | subvariant of Delta (de-escalated VOC as per the ECDC) | https://www.ecdc.europa.eu/en/covid-19/variants-concern |
| BA.2 | 4.94E-55 | | de-escalated VOC as per the ECDC | https://www.ecdc.europa.eu/en/covid-19/variants-concern |
| BA.2.10.1 | 0.005404522 | | | |
| BF.7 | 2.41E-12 | | enhanced neutralization resistance respective of parental BA.4/5 subvariant | Qu. P et al., 2023 |
| BF.7.14 | 6.84E-31 | | | |
| BN.1.2 | 1.97E-15 | | | |
| BN.1.3 | 2.69E-16 | | | |
| BQ.1 | 5.46E-14 | VOI (20-10-2022) | VOI as per ECDC (20-10-2022); enhanced neutralization resistance respective of parental BA.5 subvariant | Qu. P et al., 2023; https://www.ecdc.europa.eu/en/publications-data/spread-sars-cov-2-omicron-variant-sub-lineage-bq1-eueea |
| BQ.1.1 | 3.25E-25 | | resistant to all clinical mAbs / enhanced neutralization resistance respective of parental BA.5 subvariant | Arora P., et al. 2022; Qu P., et al., 2023 |
| BR.2.1 | 6.67E-159 | | | |
| CH.1.1 | 0.026651959 | VUM (08-02-2023) | WHO variant under monitoring since (08-02-2023) | https://www.who.int/activities/tracking-SARS-CoV-2-variants |
| CH.1.1.1 | 0.000159952 | | | |
| DY.2 | 5.38E-08 | | | |
| EG.1 | 2.85E-08 | | | |
| EG.2 | 0.00019372 | | | |
| EL.1 | 0.012517337 | | | |
| FE.1 | 8.68E-07 | VUM | VUM as per ECDC | https://www.ecdc.europa.eu/en/covid-19/variants-concern |
| FE.1.2 | 4.23E-25 | | | |
| FL.2 | 1.08E-22 | | | |
| FR.1 | 0.020160177 | | | |
| Unassigned | 0.029477699 | | | |
| XBB | 1.42E-58 | VUM (12-10-2022) | XBB and its sublineages have been identified as variants under monitoring as per the WHO (12-10-2022) | https://www.who.int/activities/tracking-SARS-CoV-2-variants |
| XBB.1 | 1.60E-05 | | | |
| XBB.1.15 | 0.002277383 | VOI (11-01-2023) | WHO current variant of interest (as of 11-01-2023) | https://www.who.int/activities/tracking-SARS-CoV-2-variants |
| XBB.1.16 | 5.97E-10 | VOI (17-04-2023) | WHO current variant of interest (as of 17-04-2023) | https://www.who.int/activities/tracking-SARS-CoV-2-variants |

| | | | | |
|---|---|---|---|---|
| XBB.1.16.1 | 0.006501881 | | | |
| XBB.1.18.1 | 2.35E-22 | | | |
| XBB.1.5 | 4.52E-22 | | | |
| XBB.1.5.12 | 2.84E-05 | | | |
| XBB.1.5.25 | 3.96E-41 | | | |
| XBB.1.5.33 | 5.67E-05 | | | |
| XBB.1.9.1 | 3.02E-05 | VUM (30-03-2023) | WHO variant under monitoring since (30-03-2023) | https://www.who.int/activities/tracking-SARS-CoV-2-variants |
| XBB.1.9.2 | 2.18E-07 | VUM (26-04-2023) | WHO variant under monitoring since (26-04-2023) | https://www.who.int/activities/tracking-SARS-CoV-2-variants |
| XBB.2 | 8.09E-58 | | | |
| XBB.2.3.2 | 6.99E-10 | | | |
| XBB.2.6 | 1.24E-115 | | | |
| XBF | 2.68E-08 | | increased antibody neutralization | Ackerman A., et al. 2023 |
